# A remarkable psyllomorph family from Cretaceous Burmese amber, Miralidae stat. nov. (= Dinglidae syn. nov.; Hemiptera: Sternorrhyncha)

**DOI:** 10.1101/2024.10.04.616632

**Authors:** Grigory A. Ivanov, Dmitry D. Vorontsov, Dmitry E. Shcherbakov

**Affiliations:** Borissiak Paleontological Institute, Russian Academy of Sciences, Moscow 117647, Russia; Koltzov Institute of Developmental Biology, Russian Academy of Sciences, Moscow 119334, Russia

**Keywords:** Hemiptera, Psyllomorpha, new taxon, Burmese amber, labium, wax pores

## Abstract

Cretaceous resins have preserved a remarkable diversity of sternorrhynchans. Burmese amber, formed in the mid-Cretaceous tropics, contains more numerous and diverse Psyllomorpha than all other fossil localities of the period together. The genus *Dingla* Szwedo et Drohojowska, 2020, previously placed into a separate family and infraorder, turned out to be similar in all essential characters to *Mirala* Burckhardt et Poinar, 2019 and *Burmala* Liu et al., 2021; therefore we synonymize Dinglidae Szwedo et Drohojowska, 2020 with Miralinae Shcherbakov, 2020 and raise the last subfamily to the full family status. A sister-group relationship between Dinglomorpha and Aleyrodomorpha in the cladistic analysis of Drohojowska et al. (2020a) was based on incorrect interpretation of characters. *@@ala @@ioides* gen. et sp. nov. from Burmese amber differs from other miralid genera by spotted forewings without pterostigma and the structure of terminalia in both sexes. In Miralidae, we discovered a long annulated labium and compound wax pores on the ventral side of the abdomen—for the first time among Hemiptera. The compound wax pores may have helped prevent miralids from sticking to the honeydew excreted. The long, flexible second labial segment, reinforced by sclerotized rings, probably allowed the protrusible length of the stylet bundle to be increased by arching the labium, as in mosquitoes.

## 1. Introduction

Burmese amber, mined in the Hukawng Valley, Kachin State, Myanmar, is a genuine treasure trove of fossils. This is not the only but certainly the richest amber deposit in Gondwana, rivaled only by Lebanese amber. Burmese amber has preserved a vast assemblage of Mesozoic organisms, including a megadiverse hexapod fauna with nearly 500 families, over 1,300 genera, and over 2,000 species described to date (Ross, 2024). The amber-containing rock had been dated to the earliest Cenomanian (ca. 99 Ma; Shi et al., 2012), although an Albian ammonite was discovered above the amber layer (Cruickshank and Ko, 2003). The amber was formed at low paleolatitude (about 12°N) in the mid-Cretaceous tropics on an isolated terrain in the Tethys Ocean (Westerweel et al., 2019). The source tree of the amber was similar to *Agathis* (Araucariaceae) and grew in a tropical-subtropical rainforest resembling the kauri forest of northern New Zealand (Poinar et al., 2007).

Some authors have suggested that the environment of Myanmar forests had not changed since the Cretaceous, as many taxa found in Burmese amber still exist today in Myanmar and other tropical and subtropical regions (Johnson et al., 2022). However, the Burmese amber flora differs strikingly from the present-day floras in the low diversity and rarity of angiosperms, which dominate modern plant assemblages (Poinar, 2022). Accordingly, the insect pollinators and specialized herbivores, the latter including many hemipterans, of Burmese amber are very different from such insects inhabiting the modern tropics, where they are mainly associated with angiosperms. For example, the psyllomorphs and aphids of Burmese amber are represented only by extinct families. The subject of this paper is one unusual psyllomorph family endemic of Burmese amber.

One of the four major extant groups of Hemiptera Sternorrhyncha, jumping plant lice or psyllids (Psylloidea s.str.), represent the crown part of an ancient phyletic lineage, the infraorder Psyllomorpha, which can be traced back to the Middle Permian (270 Ma; Becker-Migdisova, 1985). The earliest Psylloidea s.str. are recorded in the Eocene (about 45 Ma; Klimaszewski, 1997; Ouvrard et al., 2013). Two Mesozoic families, Liadopsyllidae and Malmopsyllidae, were united with psyllids under Psylloidea s.l. (Burckhardt and Poinar, 2019; Shcherbakov, 2020). Permian and Mesozoic Protopsyllidiidae and their allies constitute another superfamily.

Modern psyllids are in many respects much more derived than their extinct precursors. The labium in psyllids is barely capable of moving, short, stout, much shorter than the stylet bundle contained in it. When not feeding, the excess length of the bundle is looped in the pocket (crumena) inside the thorax in adults, or outside in front of the labium in nymphs; when the stylets penetrate into the host plant, the stylet loop straightens out (Weber, 1929, 1933). In the known extinct psyllomorphs, the labium is long and slender, while the crumena and the stylet loop have never been recorded.

Psyllids jump with their hind legs, like Auchenorrhyncha and some true bugs, but their jumping mechanism is very peculiar. They use their hind legs, with highly modified coxae, to take off with a series of rapid forward flips, creating an unpredictable trajectory, often without switching to flight (Burrows, 2012). The hind coxae are unmodified in modern “half-jumping” psyllids (Togepsyllinae), probably due to secondary reduction (Luo et al., 2017), and in the extinct psyllomorph families.

Like other Sternorrhyncha, psyllids feed on phloem and their liquid excrements (honeydew) are rich in sugar (Douglas, 2006). This creates the risk of the insect getting stuck to the contaminated surface of the host plant. Many aphids are attended by ants, which both protect them and remove the honeydew. Among psyllids, the ant attendance is a rare exception (Shrestha et al., 2022). Instead, in psyllid nymphs and females, the circumanal ring of wax glands packs honeydew into bags of wax film, while the males use their mobile proctiger to place sticky honeydew droplets closely behind their body (Ammar et al., 2013a).

The highest diversity of Mesozoic psyllomorphs has been recorded from the Jurassic (Li et al., 2022). The richest Cretaceous psyllomorph fauna is found in mid-Cretaceous Burmese amber. Besides protopsyllidioids and typical liadopsyllids, a peculiar psyllomorph, *Mirala burmanica* Burkchardt et Poinar, 2019, has been described from Burmese amber and assigned to Liadopsyllidae s.l. (including Malmopsyllidae). Next year, the genus *Mirala* was placed into the subfamily of its own, Miralinae, within the resurrected Malmopsyllidae (Shcherbakov, 2020). Later the same year, *Dingla shagria* Szwedo et Drohojowska, 2020 was described as representing a separate family, Dinglidae, and even a new infraorder, Dinglomorpha, considered to be a sister group to Aleyrodomorpha (Drohojowska et al., 2020a). Later on, *Alloeopterus anomeotarsus* Poinar et Brown, 2020 was described in Dinglidae and *Burmala liaoyaoi* Liu et al., 2021 in Miralinae.

Based on the study of original descriptions and new material, we came to the conclusion that the genera *Mirala* Burckhardt et Poinar, 2019, *Burmala* Liu et al., 2021, and *Dingla* Szwedo et Drohojowska, 2020 were similar in all main characters (see 4.1) and should be placed in one family, Miralidae stat. nov. (= Dinglidae syn. nov.) within Psylloidea s.l. More data are needed to clarify the systematic position of *Alloeopterus* Poinar et Brown, 2020. A new miralid genus and species is described below, *@@ala @@ioides* gen et sp. nov. The separation of this family into an infraorder of its own is not justified, although such characters of Miralidae as the long annulated labium and compound wax pores on the ventral side of the abdomen have not previously been recorded in Hemiptera.

## 2. Material and methods

The Burmese amber is collected from the mid-Cretaceous deposits of Hukawng Valley, northern Myanmar. The amber mines are located at the north end of Noije Bum hill about 20 km SW of Tanai. The horizon of the amber mining is upper Albian to lower Cenomanian; a radioisotopic date has been established at 98.79 ± 0.62 Ma (Shi et al., 2012). A detailed locality map and stratigraphic information are given by Cruickshank and Ko (2003).

One male and one female of *@@ala @@ioides* gen et sp. nov., as well as five specimens of *Dingla shagria* and one of *Burmala liaoyaoi* were found in pieces of Burmese amber purchased on ebay.com in 2017. Now the pieces are in the collection of the Paleontological Institute, Russian Academy of Sciences (PIN).

Pieces with inclusions were carefully processed using a hand-held rotary saw Proxxon LU-6868 and a polishing machine (Sidorchuk and Vorontsov, 2018). In this way, processed amber fragments were obtained, which can be photographed under a microscope from any side (Fig. 1B).

**Fig. 1.**
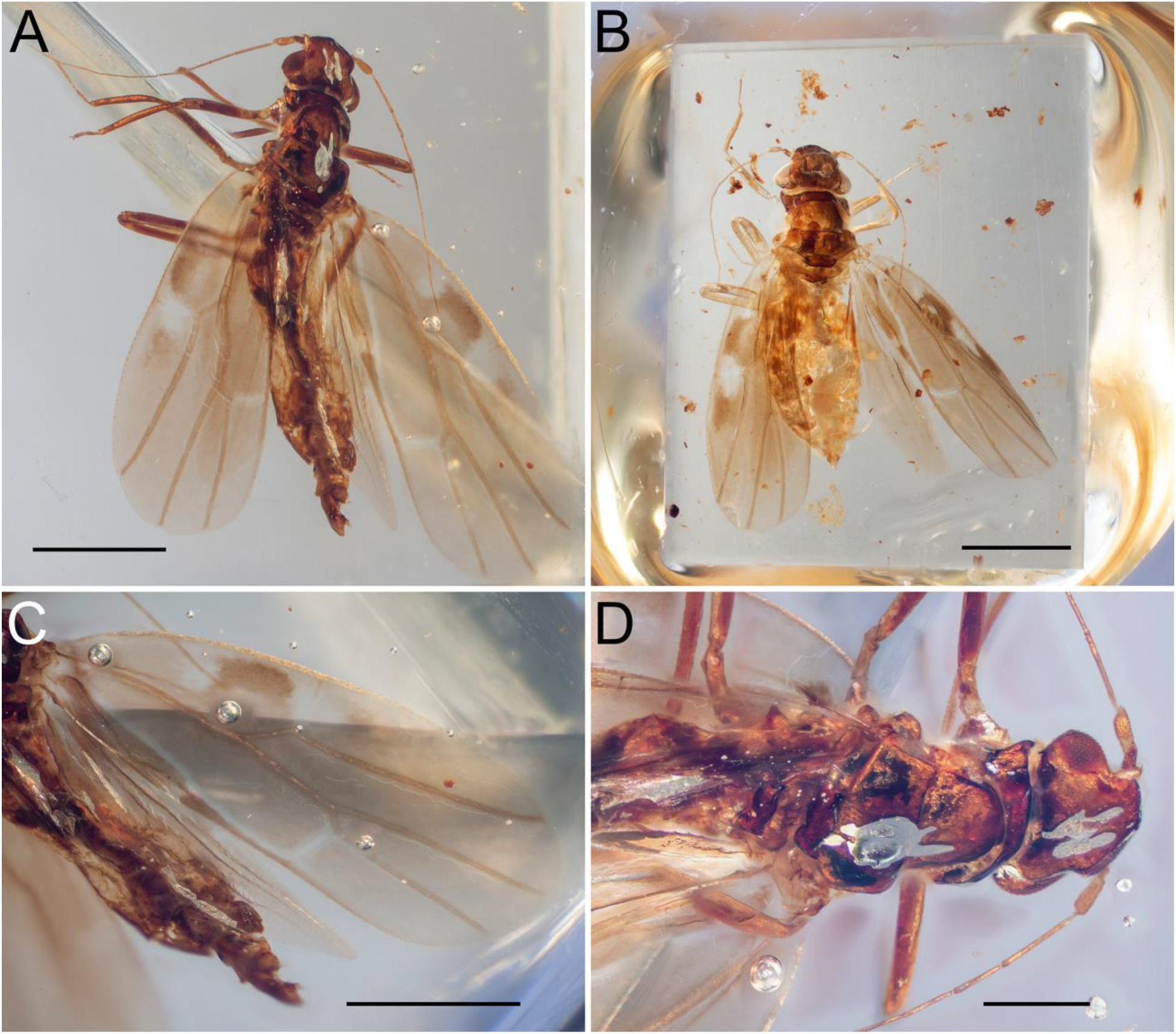
*@@ala @@ioides* gen. et sp. nov.: holotype male (A, C, D) and paratype female (B): (A) male, habitus; (B) female, habitus; (C) fore and hind wing, dorsal view; (D) forebody, dorsal view. Scale bars: 0.5 mm (A–C); 0.2 mm (D).

**Fig. 2.**
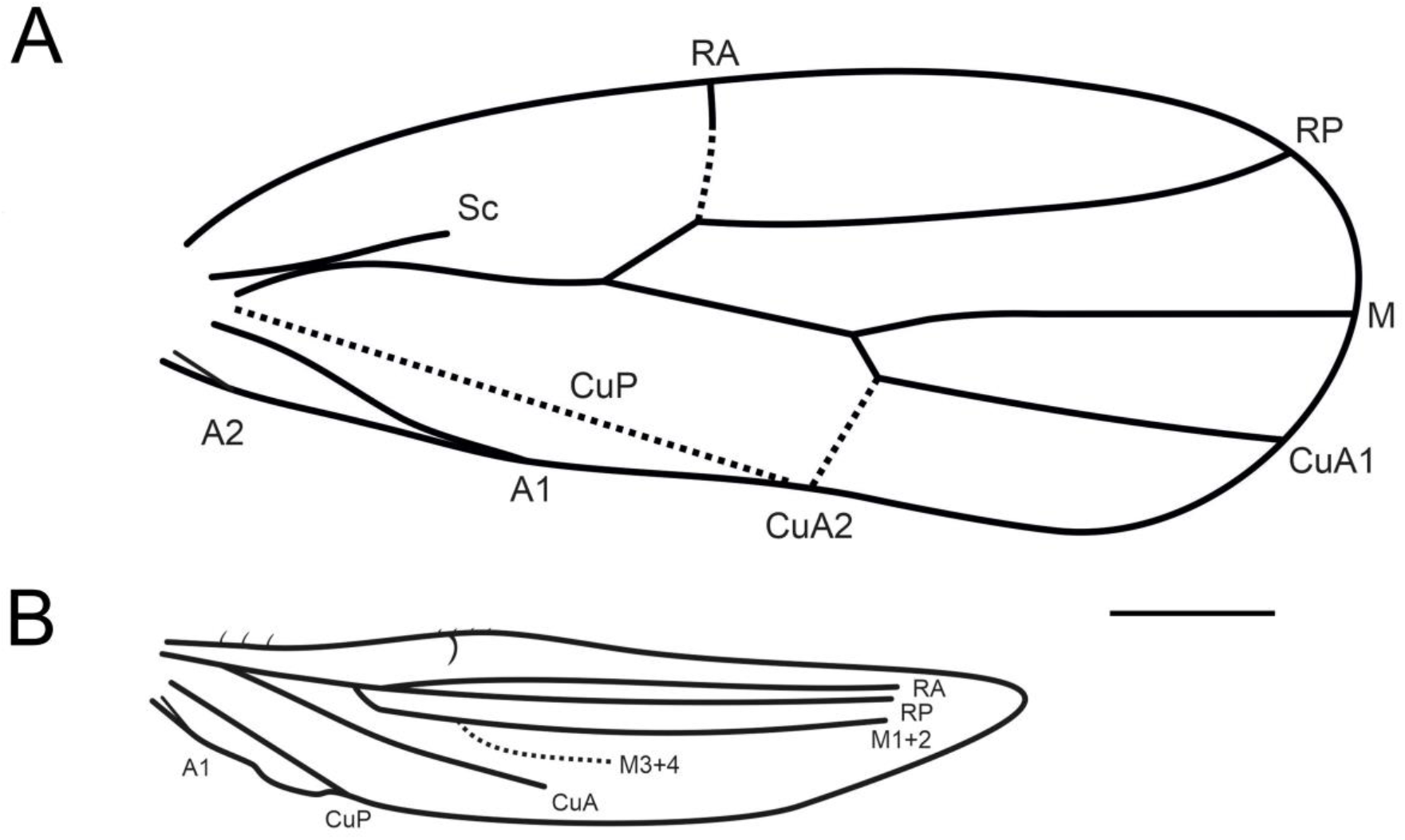
*@@ala @@ioides* gen. et sp. nov., venation, holotype male: (A) forewing; (B) hind wing; dotted line, fold-like or weak veins. Scale bar: 0.3 mm.

**Fig. 3.**
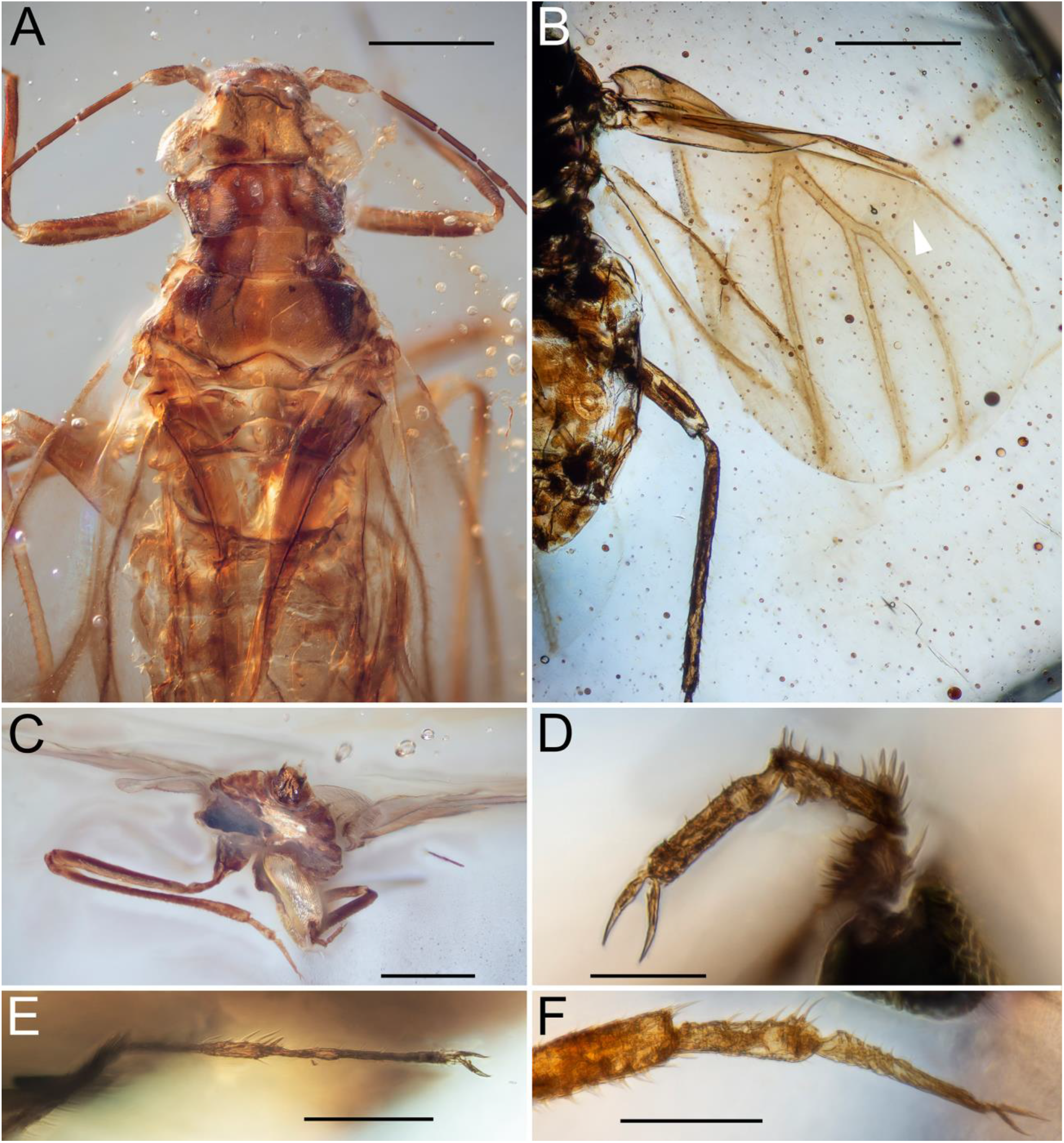
*Dingla shagria* Szwedo et Drohojowska, 2020, males, dorsal view (A–B): (A) PIN 5608/212a, forebody; (B) PIN 5608/177a, wings (arrowhead, forewing CuA2). *@@ala @@ioides* gen. et sp. nov., holotype male (C–F): (C) caudal view; (D–F) tarsus and apex of tibia of right (D, E) and left (F) hind leg (proportions distorted). Scale bars: 0.3 mm (A–C); 0.05 mm (D); 0.1 mm (E, F).

For imaging, a Nikon E-800 compound microscope with dry (4x and 10x) and water immersion (40x and 60x) objectives was used, equipped with an Olympus OM-D E-M10 II digital camera (Sidorchuk and Vorontsov, 2018), as well as stereomicroscopes Nikon SMZ25 with Nikon DS-Ri2 digital camera and Leica M165C with Leica DFC425 digital camera, to obtain image stacks representing multiple focal planes. In addition, we used a Zeiss LSM 880 confocal microscope and a Leica Thunder DMi8 fluorescent microscope. The resultant images were focus-stacked in Helicon Focus Pro 7.0.2.

We also used a Tescan Vega microscope to obtain scanned electron micrographs of uncoated modern psyllids from the collection of the Zoological Institute RAS, St. Petersburg (ZIN).

## 3. Systematic paleontology

Order Hemiptera Linnaeus, 1758

Suborder Sternorrhyncha Amyot et Serville, 1843

Infraorder Psyllomorpha Becker-Migdisova, 1962

Superfamily Psylloidea Latreille, 1807, sensu lato

Family Miralidae Shcherbakov, 2020, stat. nov. = Dinglidae Szwedo et Drohojowska, 2020, syn. nov.

Miralidae: Shcherbakov, 2020 (June 26)

Dinglidae: Drohojowska et al., 2020 (July 9), syn. nov.

### Diagnosis

Body dorsoventrally flattened. Forewing with or without pterostigma; M unforked; CuA2 recurrent, sometimes desclerotized; apex of clavus about wing midlength; CuA1 straight, directed towards apical wing margin; R bifurcation at 0.3–wing length, M+CuA bifurcation at 0.6–0.7 wing length. Head with compound eyes undivided; three ocelli present, median ocellus located frontally; antennae long, 10-segmented, filiform; subapical rhinarium may be present on each flagellomere; one (subapical or apical) or no seta on last flagellomere. Labium long, reaching at least to metacoxae; 4-segmented: 1st segment short, hidden between prothoracic lobes; 2nd segment longest, with 50–80 narrow sclerotized rings; 4th shortest, directed ventrally. Thorax with pentagonal or subtrapezoidal meso- and metascutellum. Hind legs longest; coxae not modified; tarsi 2-segmented, without empodium, arolium or pulvilli. Abdomen with three pairs of compound wax pores, each with ring-shaped sclerite, on ventral laterotergites 3–5 (male) or 4–6 (female). Male terminalia either with short subgenital plate and long parameres or with long subgenital plate and short parameres; each paramere armed with hooked claw and apical process with thick setae. Female terminalia with very short or relatively long proctiger, outer valvulae shaped similar to or different from inner valvulae, and subtriangular subgenital plate.

### Composition

*Mirala* Burckhardt et Poinar, 2019; *Dingla* Szwedo et Drohojowska, 2020; *Burmala* Liu et al., 2021; *@@ala* gen. nov.

### Distribution

Mid-Cretaceous amber of Myanmar.

### Remarks

The monotypic genus *Alloeopterus* Poinar et Brown, 2020 from Burmese amber (originally described in Dinglidae) is similar to Miralidae in the dorsoventrally flattened body and unforked M in the forewing, but differs from this family in the proximal R fork, oblique RA, reduced CuA fork in the forewing and the non-annulated labium. More data are needed to clarify the position of this genus within Psylloidea.

Genus *@@ala* gen. nov.

(urn:lsid:zoobank.org:

Type species: *@@ala @@ioides* gen. et sp. nov.; by original designation and monotypy.

### Etymology

From @@@@@@@@@. Gender: feminine.

### Diagnosis

Forewing with conspicuous dark and pale pattern; pterostigma absent; RA and CuA2 short, transverse, desclerotized; free CuA stem very short; A1 sigmoidal. Antenna longer than head+thorax; flagellum very slender; last flagellomere with one subapical rhinaria and without terminal concavity. Labium reaching metacoxae; second labium segment with about 50 sclerotized rings. Pronotum short, arched; meso- and metascutellum slightly W-shaped. Metatibia with several thick apical setae. Parameres short, each with thick apical setae; proctiger short, with apical anus; male subgenital plate much longer than proctiger. Female genitalia partially covered by subgenital plate from below.

### Remarks

The new genus is distinct from *Burmala* and similar to other genera in the rostrum reaching only to the metacoxae. It also differs from all genera of the family in its spotted forewings without pterostigma, long and slender antennae, a short pronotum, a long subgenital plate, and short proctiger and parameres.

*@@ala @@ioides* gen. et sp. nov.

(urn:lsid:zoobank.org:

Figs 1–8

### Etymology

@@@@@@@.

### Type material

Holotype, male: PIN 5608/330; Paratype, female: PIN 5608/214

### Locality and horizon

Mid-Cretaceous, Tanai Village, Hukawng Valley, Kachin State, northern Myanmar.

### Diagnosis

As for genus, with some additional characters: Metatibia with rows of thick, erect, curved setae along outer side and at least 5 thick apical setae. Metabasitarsus with several thick setae on plantar side. Male subgenital plate scoop-shaped; each paramere with darkened sclerotized basal claw and long, paired thick apical setae.

### Description

Total length and width (including wings as preserved) 1.5 × 1.6 mm in female and 1.8 × 2.1 in male.

### Head

Compound eyes large, elongated, undivided, of similar ommatidia. Three ocelli; lateral ocelli close to anterior margin of eyes; median ocellus mostly visible from dorsal side (Fig. 4A). Antennae 10-segmented, filiform; pedicel 1.5x longer than scape; flagellum thin, 3x narrower than pedicel; antennomeres 3, 9, 10 of the same length, 2x longer than antennomeres 4–8. Antennomeres 3–8 with annular pattern, covered with minute hairs and bearing subapical setae (Fig. 4C); antennomeres 9 and 10 densely transversely ridged, bare, except subapical setae on antennomere 9 and one apical seta on antennomere 10 (Fig. 4D–F); subapical rhinaria on antennomeres 3–10 (biggest on 3, 5) in both sexes; apical antennomere without processus terminalis or terminal concavity, bears 1 apical seta on a slightly separated rounded apex, and is covered with small areas of short denticles (Fig. 4D–F). Labium long (Fig. 5A), reaching metacoxae; 4-segmented: 1st segment short, hidden between prothoracic lobes, 2nd segment very long, bears about 50 regular sclerotized narrow annuli and regularly placed setae (every third or fourth annulus; Fig. 5B, C), 3rd 5–6x shorter than 2nd, 4th directed perpendicular to 3rd, 2–2.5x shorter than 3rd (Fig. 5B); in distal portion wider. *Thorax*. Pronotum arched, short, 7x wider than long; mesoprescutum large, as long as wide, almost not covered by pronotum; mesoscutum thrice wider than long; mesoscutellum and metascutellum slightly W-shaped (Fig. 1D).

**Fig. 4.**
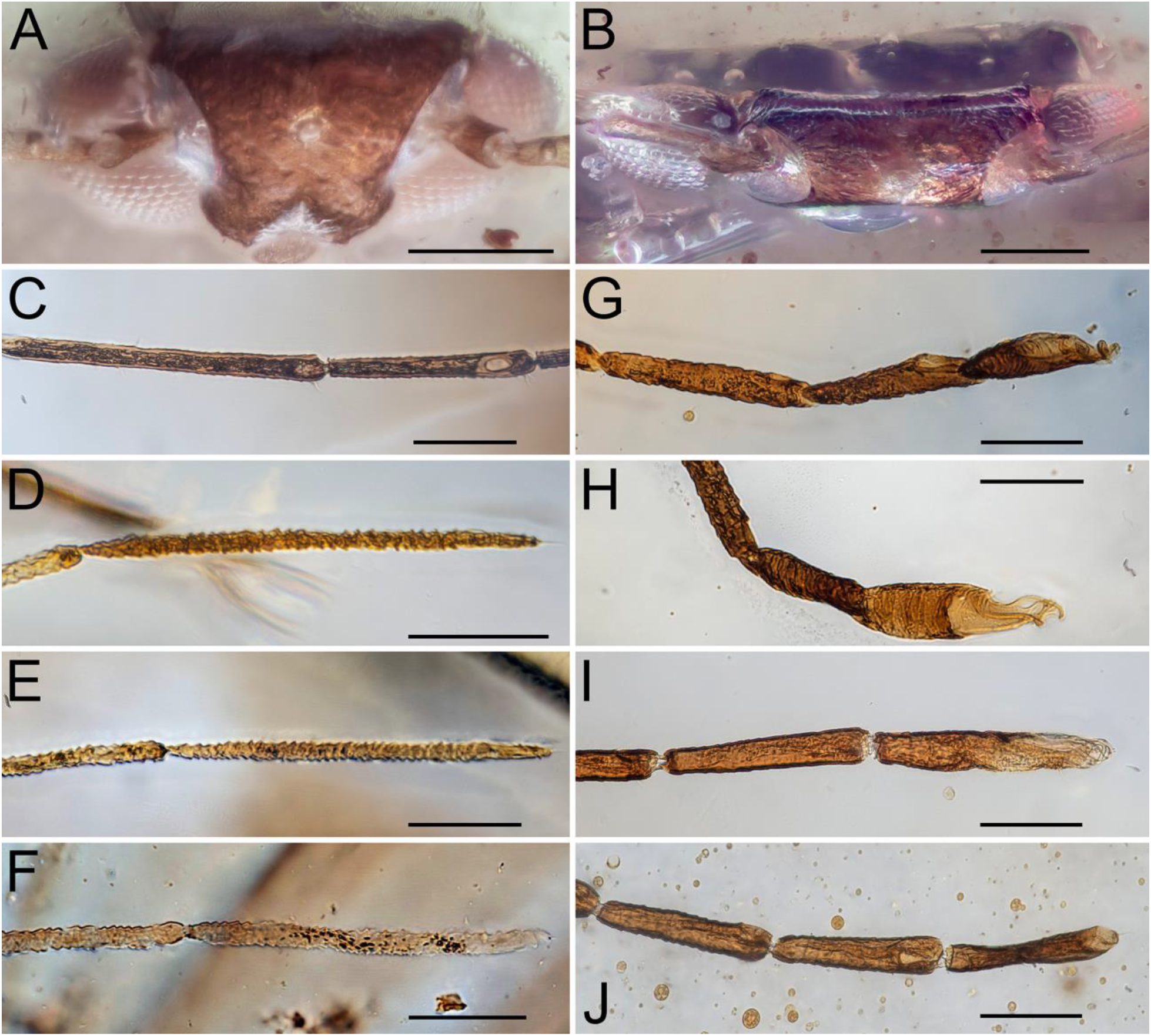
Head, anterior view (A, B): (A) *@@ala @@ioides* gen. et sp. nov., paratype female; (B) *Dingla shagria*, female PIN 5608/213. Antennae (C–J): subapical rhinaria on flagellomeres 1, 2 (C); distal flagellomeres (D–J): *@@ala @@ioides* gen. et sp. nov., holotype male (C–E) and paratype female (F); *Dingla shagria*, female PIN 5608/213 (J) and male PIN 5608/212a (I); *Burmala liaoyaoi* Liu et al, 2021, male PIN 5608/122a (G, H). Scale bars: 0.1 mm (A–B); 0.05 mm (C-J).

**Fig. 5.**
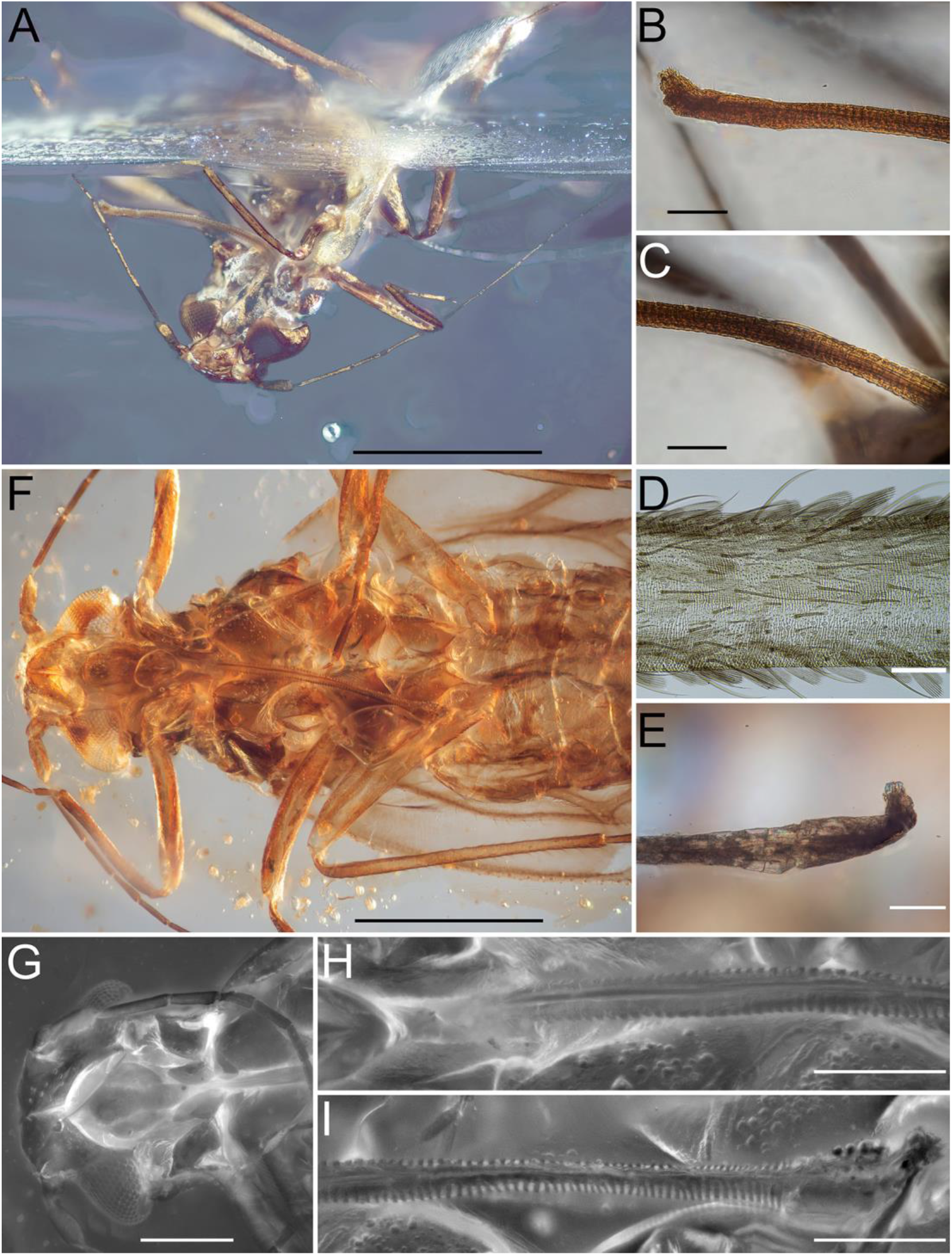
Annulated labium of Miralidae (A–C, E–I) and Culicidae (D): (A–C) *@@ala @@ioides* gen. et sp. nov., holotype male: (A) forebody, anteroventral view; (B–C) distal (B) and proximal (C) parts of labium; (D) recent Culicidae gen. sp., part of prementum with alternating sclerotized and membranous stripes of cuticle (photo by R. Rakitov); (E) *Burmala liaoyaoi*, male PIN 5608/122a, 3rd and 4th labial segments; (F–I) *Dingla shagria*, males PIN 5608/212a (F, H–I) and PIN 5608/177a (G), ventral view: (F) forebody; (G) head and prothorax; (H–I) proximal (H) and distal (I) parts of labium; (G–I) under fluorescence. Scale bars: 0.5 mm (A, B); 0.05 mm (C–F); 0.2 mm (G); 0.1 mm (H, I).

### Legs

Coxae and trochanters very long (coxa+trochanter 1/2 femur length; Figs 1A, D, 3C); metacoxa without meracanthus. Protibia and profemur similar in length, mesotibia and mesofemur similar in length, metatibia 1.5x longer than metafemur; protibia with row of thick setae on outer side and 2 thick apical setae (Fig. 5A); mesotibia with rows of setae on outer side (Fig. 5A); metatibia widened distally with rows of thick erect, curved setae on outer side (turning thicker distally) and at least 5 thick apical setae (Fig. 3C). Tarsi 2-segmented; protarsus 1.5x as long as protibia, mesotarsus 3x as long as mesotibia, metatarsus 2.2x as long as metatibia; probasitarsus 2x as long as prodistitarsus; mesobasitarsus 1.5x as long as mesodistitarsus; metabasitarsus 0.9x as long as metadistitarsus; metabasitarsus with row of six thick setae on plantar side; claws with short tooth near base; pretarsus with no arolium, empodium, or pulvilli (Fig. 3D–F).

### Forewing

Forewing 2.5x as long as wide, with conspicuous dark and pale pattern: darkened areas – on distal part of costal space and slightly beyond nodal line; pale areas on basal part of costal space and on nodal line (Fig. 1A–C). Pterostigma absent; bSc undeveloped. RA and CuA2 short, transverse, desclerotized; M unforked; free CuA stem very short (Figs 1A, C, 2A). R bifurcation at 0.4 wing length, M+CuA bifurcation at wing length. A1 wavy (Fig. 1C). Claval furrow distinct; claval apex about 1/2 wing length; claval vein reaching posterior margin (Figs 1C, 2A). Costal and apical margins bearing short setae; veins with sparse short setae. Coupling fold with row of small denticles on claval margin (Fig. 1C).

### Hind wing

Hind wing 0.7x as long and 0.5x as wide as forewing; costal margin with single strong coupling hook, proximal row of dorsally directed setae and distal row of short denticles (Figs 1A, C, 2B); C sclerotized proximally and weak distal to hook (Fig. 1C); M unforked in paratype female (Fig. 1B) and forked in holotype male (Figs 1C, 2B), M3+4 short and weak; RA, RP, M parallel distally, weakened apically, not reaching wing margin; CuA reaching wing midlength; anal vein reaching posterior margin (Figs 1C, 2B).

### Female abdomen

Tergites 3–8 and proctiger distinct; tergite 8 narrow; proctiger small with large oval anus; sternites 3–9 distinct; sternite 9 forming small triangular subgenital plate (Fig. 6F); with 3 pairs of compound wax pores, each with ring-shaped sclerite, ventrally on laterotergites 4–6, pores of last pair 1.5x larger than others (Fig. 8C). Ovipositor short, pale, with two pairs of valvulae visible: inner valvulae slender; outer valvulae robust, with lateral setae and each with one apical spine, proximally with several folds, converging dorsolaterally, covering inner valvulae only laterally (Fig. 6E– H).

**Fig. 6.**
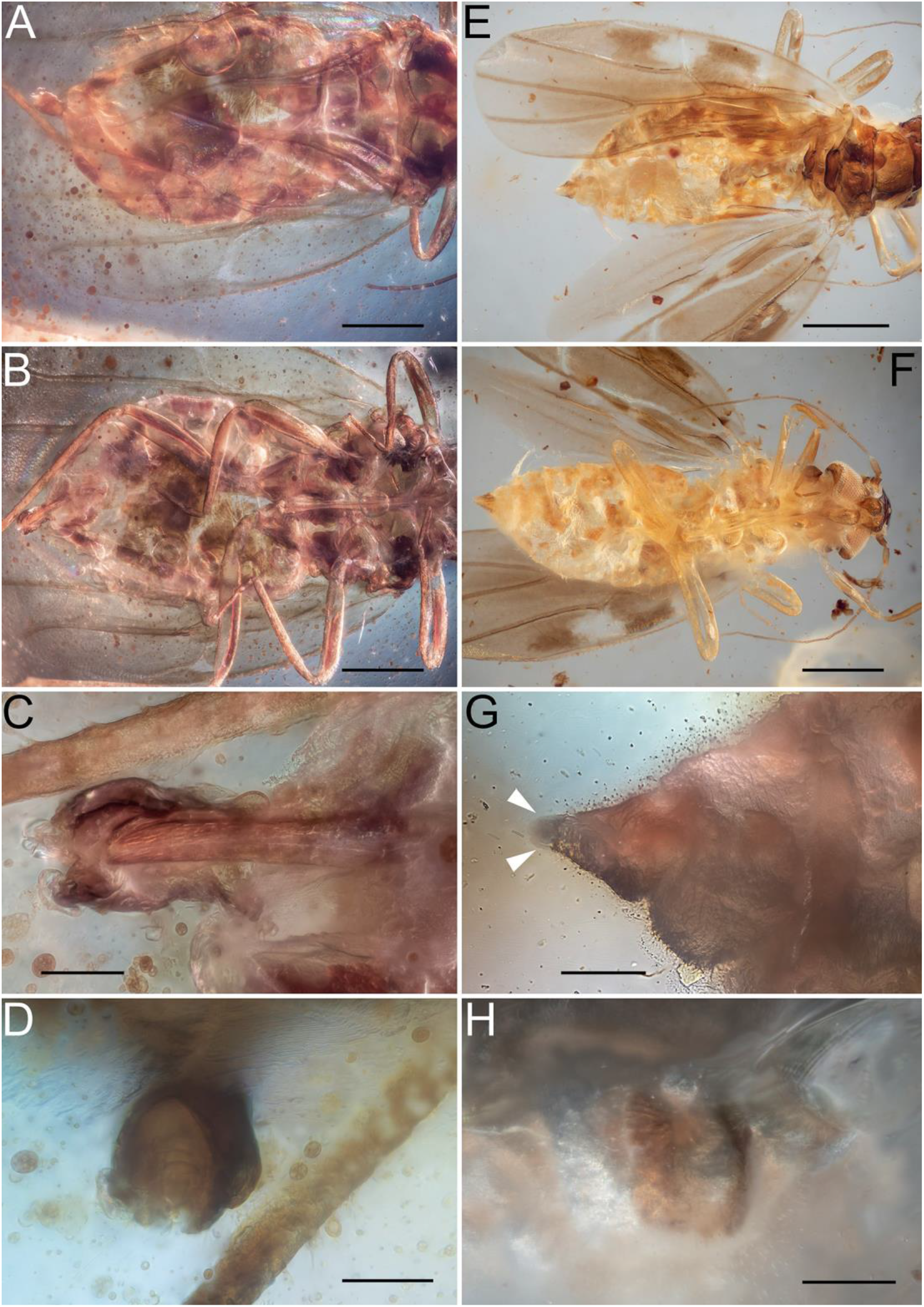
Female terminalia of Miralidae: (A–D) *Dingla shagria*, PIN 5608/213; (E–H) *@@ala @@ioides* gen. et sp. nov., paratype; (A, E, G) dorsal view; (B, C, F) ventral view; (D, H) caudal view; arrowheads, thick apical setae of outer valvulae (G). Scale bars: 0.3 mm (A, B, E, F); 0.05 mm (C, D, G, H).

### Male abdomen

Tergites 4–8 and proctiger distinct; tergite 7 0.7x as wide as tergite 6; tergite 8 shaped as narrow strip; proctiger with apical anus and several thick setae at apex; tergite 8 and proctiger slightly retracted into tergite 7; tergite 7 with lateral lobes covering subgenital plate; sternites 3–9 distinct; wax pores not found; sternite 8 with pair of triple furrows sublaterally (Fig. 1A, C); sternite 9 shaped as long, scoop-like subgenital plate with irregular sclerotized furrows and long sparse setae (Fig. 7A, B). Parameres very short, directed caudally and slightly dorsomedially; each paramere with sclerotized basal claw and apical process with one thin hair-like subapical seta and two long thick apical setae (Fig. 7A, B). Aedeagus indistinguishable.

**Fig. 7.**
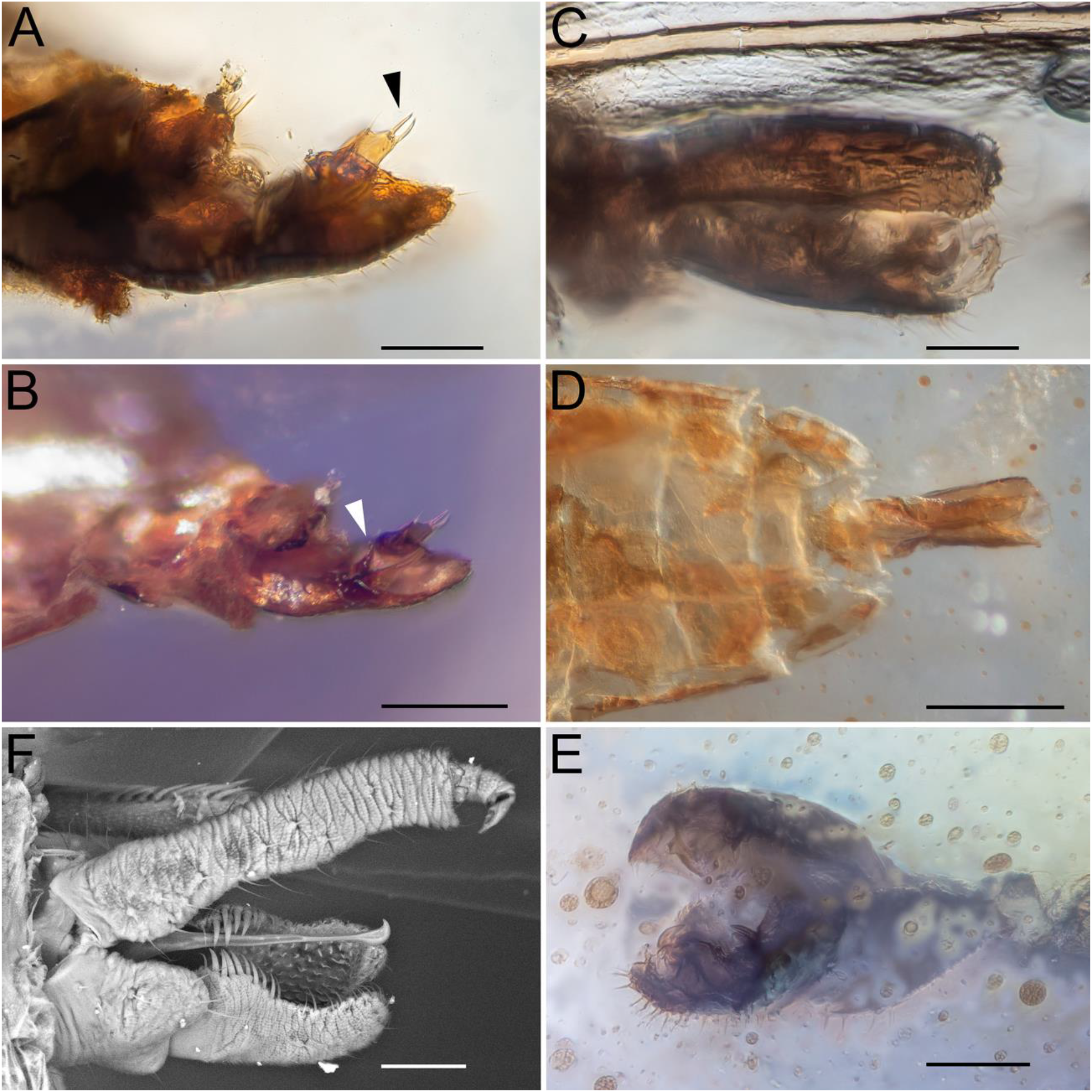
Male terminalia of Miralidae (A–E) and Togepsyllinae (F): (A–B) *@@ala @@ioides* gen. et sp. nov., holotype, lateral view; (C–E) *Dingla shagria*: (C) PIN 5608/212a, dorsolateral view; (D–E) PIN 5608/288, ventral (D) and caudolateral (E) view; (F) *Togepsylla matsumurana* Kuwayama, 1949, SEM, ZIN; arrowheads, thick setae on apical process (A) and basal claw (B) of parameres. Scale bars: 0.05 mm (A, C, E, F); 0.1 mm (B); 0.2 mm (D).

**Fig. 8.**
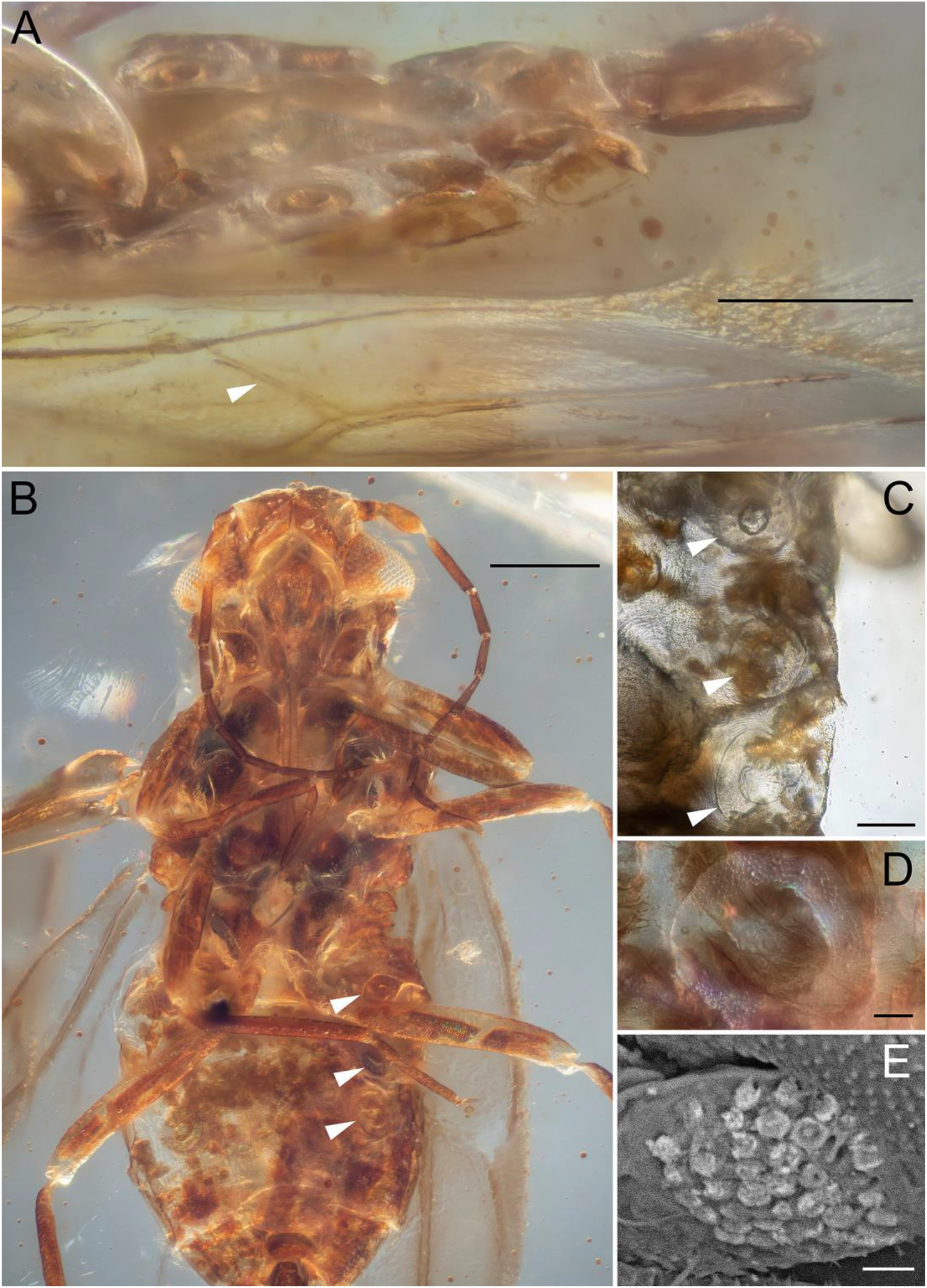
Compound wax pores of Miralidae (A–D) and wax field of Togepsyllinae (E): (A, B) *Dingla shagria*: (A) male PIN 5608/288, abdomen and forewing, ventrolateral view; (B) male PIN 5608/177a, ventral view; (C) *@@ala @@ioides* gen. et sp. nov., paratype female, abdomen, ventral view; (D) *Dingla shagria*, female PIN 5608/213, compound wax pore of 5th abdominal segment; (E) *Togepsylla matsumurana*, male, wax pore field of 5th abdominal segment, SEM, ZIN; arrowheads, forewing CuA2 (A) and abdominal compound wax pores (B, C). Scale bars: 0.2 mm (A); 0.3 mm (B); 0.05 mm (C); 0.025 mm (D); 0.005 mm (E).

### *Measurements* (mm)

Total length: 1.5 in female, 1.8 in male. Forewing length: 1.25 in female, 1.35 in male; forewing width: 0.5 in female, 0.6 in male. Antennomere length: scape 0.05–0.06 in female, 0.07–0.08 in male; pedicel 0.07–0.08 in female, 0.1 in male; antennomeres 3, 9, 10 about 0.15 in female, about 0.2 in male; antennomeres 4–8 about 0.07 in female, about 0.1 in male. Labium length: 0.41 in female, 0.47 in male. Tarsomere length: probasitarsus and prodistitarsus: 0.04 and 0.1 in female; 0.075 and 0.14 in male; mesobasitarsus and mesodistitarsus: 0.06 and 0.1 in female, 0.07 and 0.11 in male; metabasitarsus and metadistitarsus: 0.075 and about 0.09–0.11 in male, about 0.11–0.13 and about 0.1 in female.

### Additions to diagnoses of known genera

Genus *Mirala* Burckhardt et Poinar, 2019

Costal vein without break at nodus (presumed break being boundary between slender costa and thickened pterostigma in anterior view). First flagellomere as long as second. Last flagellomere gradually narrowed beyond subapical rhinarium.

*Mirala burmanica* Burckhardt et Poinar, 2019

Known only from an incompletely preserved male holotype with partly destroyed genitalia.

Genus *Dingla* Szwedo et Drohojowska, 2020

Forewing with CuA2 distinct (Figs 3B, 8A; Drohojowska et al., 2020a, figs. 2A, 3A). First flagellomere longer than second. Last flagellomere with apical group of rhinaria, one subapical (males) or one apical (females) seta, small terminal concavity, and without processus terminalis (Fig. 4I, J). Second labial segment with about 80 sclerotized annuli; 3rd 0.13–0.17 as long as 2nd, 4th directed ventrally, 0.29–0.34 as long as 3rd (Fig. 5F). Mesoscutellum well-developed (Fig. 3A). Abdomen with three pairs of compound wax pores on ventral laterotergites 3–5 (males; Fig. 8B) or 4–6 (females; Fig. 6B). Male proctiger smooth, boat-shaped (Fig. 7C). Parameres closely connected or fused, each with pale apical process bearing long lateral setae and two short thick apical setae and with sclerotized hook-like claw more medially (Fig. 7C–E). Female proctiger with apical anus, directed caudally, covering ovipositor dorsally; ovipositor tubular with subequal pairs of valvulae; subgenital plate not covering ovipositor ventrally (Fig. 6A–D).

*Dingla shagria* Szwedo et Drohojowska, 2020

New material. Specimens PIN 5608/177a, 5608/212a, 5608/213 (female), 5608/288.

Genus *Burmala* Liu et al., 2021

First flagellomere longer than second. Last flagellomere with group of rhinaria and terminal concavity, scoop-shaped, without apical or subapical setae (Fig. 4G, H). Second labial segment annulated, fourth segment with several apical lobes; 3rd ∼0.17 as long as 2nd; 4th laterally directed, 0.4–0.5 as long as 3rd (Fig. 5E). Abdomen with three pairs of compound wax pores on ventral laterotergites 3–5. Male proctiger slightly curved upward near middle, with spinous process at apex and two lateral comb-like rows of denticles along distal margin. Parameres closely connected or fused, each with lateral pale apical process and submedial setal rows.

*Burmala liaoyaoi* Liu et al., 2021

New material. Specimen PIN 5608/122a.

### Key to the genera of Miralidae

1. Forewings with extensive dark spots; pterostigma not developed. Antennae long and slender. Pronotum short ……………………………………………………………. *@@ala* gen. nov. – Forewings transparent or uniformly fuscous; most of veins and triangular pterostigma dark. Antennae less elongate …………………………………………………………………….. 2
2. Forewing tip near RP apex. 1st flagellomere as long as 2nd. Hind tarsus with 1st segment almost half as long as 2nd ………………………………… *Mirala* Burckhardt et Poinar, 2019 – Forewing tip at M apex. 1st flagellomere longer than 2nd. Hind tarsus with 1st segment almost as long as 2nd …………..………………………..………………………………….. 3
3. Forewing membrane transparent; CuA2 and RA distinct, dark. Labium reaching base of abdomen …………..………………………..………………………………….. *Burmala* Liu et al., 2021 – Forewing membrane fuscous; CuA2 and base of RA pale, membranized. Labium reaching metacoxae ……………………………………………… *Dingla* Szwedo et Drohojowska, 2020

## 4. Discussion

### 4.1. Synonymy of Miralidae and Dinglidae

The genera *Mirala* and *Burmala* (assigned to Miralinae) and the genus *Dingla* (described in Dinglidae) are similar in all essential characters: the unique forewing venation (R fork shifted to the nodal level, M+CuA fork even more distal, M unforked, CuA fork large, CuA2 recurrent, RA transverse, RA and CuA2 often desclerotized); the dorsoventrally flattened body; the last flagellomere with an apical group of rhinaria; the elongated annulated labium; the relatively long pronotum; three pairs of compound wax pores on the ventral side of the abdomen; and the peculiar male terminalia. There is no doubt that these three genera must be placed in the same family, which differs significantly from Liadopsyllidae and Malmopsyllidae in the forewing venation and body structure, including such unique features as the annulated labium and compound wax pores on the abdomen. Therefore, we propose the following synonymy: Miralidae Shcherbakov, 2020, stat. nov. = Dinglidae Szwedo et Drohojowska, 2020, syn. nov.

Drohojowska et al. (2020a) separated the family Dinglidae into the infraorder Dinglomorpha, which appeared as the sister group of Aleyrodomorpha in their cladistic analysis due to four shared derived characters. Of these characters, two were considered homoplasies and the other two were considered synapomorphies, but, according to our data, all four are based on incorrect interpretations. The processus terminalis (the narrowed part of the last flagellomere distal to the rhinarium) was considered homoplasy for the clades Dinglomorpha + Aleyrodomorpha and Aphidomorpha + Naibiomorpha. We found that, in *Dingla*, the last flagellomere bears an apical group of rhinaria and there is no processus terminalis (see 4.2). The so-called “areola postica” (i.e. CuA fork) was coded as “absent” (another homoplasy), although the reclined, weak CuA2 is clearly visible in PIN specimens (Figs 3B, 8A) and even in photographs of *Dingla shagria* types (Drohojowska et al., 2020a, figs 2A, 3A). The other two putative synapomorphies concerned thorax structure. However, the mesoscutellum of *D. shagria*, coded as “reduced, membranous,” is, in fact, well-developed and sclerotized (see the original species description and Figs 3A, 6A), and the mesopostnotum, coded as “well-developed,” but not mentioned in the species description, is not visible in PIN specimens—perhaps some part of the metascutum was mistaken for the mesopostnotum. Therefore, the alleged sister relationship between Dinglidae and Aleyrodidae and the erection of the infraorder Dinglomorpha are not justified.

### 4.2. Antennae

The structure of antennae in Miralidae is generally similar to modern psyllids. Their antennae are long, the first flagellomere is usually the longest, and rhinaria may be present subapically on each flagellomere (Drohojowska et al., 2020a). Each flagellomere bears subapical setae and its surface has an annular pattern as in modern psyllids (Kristoffersen et al., 2006; Onagbola et al., 2008; Fig. 4C).

Modern psyllids (both nymphs and adults) have two apical setae on the last flagellomere, while adult whiteflies always have one apical seta (Haupt, 1934; Kristoffersen et al., 2006). Miralidae show varying number and arrangement of setae on the last flagellomere, in some cases even between sexes of one species: one subapical seta in males of *Dingla* (Fig. 4I), one apical seta in the female of *Dingla* (Fig. 4J); no setae in males of *Burmala* (Fig. 4G, H); and one apical seta in both sexes of *@@ala* (Fig. 4D–F). In all Miralidae, the setae on the last flagellomere are considerably shorter than in the amber Liadopsyllidae and modern psyllids (Onagbola et al., 2008; Ouvrard et al., 2010; Drohojowska et al., 2020b).

The structure of the antennal apex in Miralidae is also variable. In *Dingla* and *Burmala*, the last flagellomere bears an apical group of rhinaria, more intricately shaped in *Dingla* males than in females, and in *Burmala* with a deep terminal concavity (Fig. 4G, H). Among modern psyllids, similarly grouped rhinaria are found on the 2nd flagellomere in *Bactericera femoralis* (Foerster, 1848) (Triozidae; Klimaszewski, 1964). On the other hand, in *@@ala* the last flagellomere has no terminal concavity, its subapical rhinaria do not form a group, and the surface of the last two flagellomeres is prominently ridged (only slightly so in other cases). The last flagellomere is barely if at all narrowed beyond the subapical rhinarium in *@@ala* (Fig. 4D–F) and only gradually narrowed in *Mirala* (Burckhardt and Poinar, 2019, fig. 2b).

### 4.3. Labium

In modern psyllids the labium is directed ventrally and is about as long as the head, whereas in Miralidae the labium is greatly elongated, directed caudally at rest, reaching the metacoxae, about 0.3–0.4x as long as the body and thrice longer than the head (Fig. 5A, F). The labium is also elongated and directed caudally in *Liadopsylla apedetica* from Lower Cretaceous Lebanese amber (Ouvrard et al., 2010) and some Jurassic *Liadopsylla* species (Becker-Migdisova, 1985).

The labium of Miralidae is unique in that it is annulated for most of its length. Such a design has not been reported in other hemipterans (several wide annulations occur in the apical labial segment of some Paraprotopsyllidiidae, Protopsyllidioidea; Hakim et al., 2021). The labium is generally four-segmented in Hemiptera (Singh, 1971), and we recognize the four segments in Miralidae. The short 1st segment is poorly visible between prothoracic lobes (Fig. 5G). The very long 2nd segment (occupying 0.7–0.75 of the labium length) comprises 50 (*@@ala*) to 80 (*Dingla*) narrow sclerotized rings separated by membranous spaces (Fig. 5A, F, H, I), and bears short setae on every third or fourth ring (Fig. 5B, C). The 3rd segment is short, and the 4th is the shortest and slightly downturned (Fig. 5F). The annulated 2nd segment was apparently flexible, since in some specimens the labium is preserved with the 2nd segment bent at its base and twisted, so that the groove and the 4th segment are directed upward or sideways rather than downward (Fig. 5A).

To increase the protrusible length of the stylets for deeper penetration, hemipterans either shift the stylet bases deeper into the body, or shorten the labium (bend or telescope it), or form a loop of the stylet bundle (Rakitov, 2022). In modern psyllids, the stylet bundle is much longer than the labium and forms a loop, which is retracted into a sac (crumena) in the thorax in adults or sticks out in front of the labium in nymphs (Weber, 1929). No such loop of the stylet bundle was reported in Miralidae or other Mesozoic psyllomorphs.

In miralids, the very long 2nd labial segment itself could apparently be shortened due to partial telescoping of its numerous sclerotized rings. This assumption is supported by: (1) the rings narrow and slightly decrease in diameter towards the apex of the 2nd segment (Fig. 5H, I); (2) the rings are somewhat tapered (Fig. 5H), allowing one ring to be partially inserted into another.

In many true bugs, when piercing deeper, some labial segments can fold back like an elbow and release the stylet bundle from the groove (Cobben, 1978). In Miralidae, their long flexible 2nd labial segment could possibly bend backward (R. Rakitov, pers. comm.), or rather sideways, since Miralidae were depressed dorsoventrally and clung tightly against host plants. Indeed, in the male *@@ala* specimen, the stylet bundle is released for some distance from the 2nd labial segment (Fig. 5C). Such back- or side-arching of the labium is unknown in other Hemiptera, but is analogous to back-arching of the labium in some Culicidae females (Diptera: Nematocera) (Weber, 1933). In mosquitoes, the long tubular prementum bears alternating sclerotized and membranous stripes, resembling the structure of the 2nd labial segment in Miralidae (Robinson, 1939; Fig. 5D).

Additonally, if miralids were able to bend their very flexible labium by intrinsic muscles, controlling its curvature and perhaps even the length of the annulated 2nd segment, then these insects could have been able to change the point of penetration, while remaining motionless and cryptic on the host plant.

### 4.4. Legs

The legs of Miralidae are long and slender, especially in *@@ala*, and the hind legs are not as modified as in modern psyllids. *@@ala* has elongated conical coxae and long trochanters in all pairs of legs (Figs. 1A, D, 3C), like some other fossil psyllomorphs (Ouvrard et al., 2010). With such long legs, raising “on tiptoes” would provide enough space for the presumed bending of the flexible 2nd labial segment in miralids.

The hind legs of *@@ala* are armed with at least five thick apical setae on the tibia and with thick setae on the plantar surface of the basitarsus (Fig. 3D–F), which are not found in *Burmala, Dingla*, and Liadopsyllidae amber fossils (Ouvrard et al., 2010; Drohojowska et al., 2020a; Liu et al., 2021). Such leg armature suggests that *@@ala* used its hind legs for jumping, although the unmodified metathorax and metacoxae indicate that its jumping ability was much less developed than in modern psyllids and not as distinctive as theirs.

### 4.5. Wings

*@@ala* is similar to other Miralidae in the strong distal shift of the main vein forks, a wide costal space, an unforked M, and a wide CuA fork with a long recurrent CuA2 in the forewing, but differs in the dark-patterned forewing with a reduced pterostigma (Fig. 1). In other miralid genera, the forewing has a conspicuous pterostigma. The loss of the M fork in the forewing is characteristic of all genera of Miralidae and of most genera of the Cretaceous Paraprotopsyllidiidae. In contrast, among Psylloidea s.str., it is rarely observed, e.g. in *Liella paucivena* (Brown et Hodkinson, 1988) (= *Paurocephala paucivena = Diclidophlebia paucivena;* Liviidae *sensu* Burckhardt et al., 2023), or, sometimes, as an aberration on one or both forewings (unpublished data, specimens from the collection of ZIN).

The wing coupling apparatus of modern psyllids consists of a coupling fold on the anal margin of the forewing and a single powerful hook (interacting with the fold during flight), supplemented by a proximal row of setae (not involved in coupling during flight), on the costal margin of the hind wing (Klimaszewski, 1993). The coupling apparatus of Miralidae differs from that of modern psyllids by the presence of an additional row of distal, dorsally directed, shortened setae on the hind wing and several rows of denticles on the coupling fold of the forewing (Fig. 1C).

The contrasting dark pattern on the *@@ala* forewings is a unique feature among Miralidae and Mesozoic psyllomorphs in general. The pattern is the same in males and females.

### 4.6. Female terminalia

Females were discovered, in the genera *@@ala* and *Dingla*, for the first time in Miralidae. The morphology of female terminalia in these genera turned out to be very different.

The *Dingla* female has a large proctiger, not narrowed distally, which forms a kind of sheath covering the rather long tubular ovipositor from above and from the sides (Fig. 6A–D). The anus is located at the apex of the proctiger and directed caudally, whereas in modern psyllids the anus is mostly placed at the base of the female proctiger and directed dorsally (with a few exceptions, such as *Bactericera daghestanica* (Gegechkori, 1980) with the anus subapical). The ovipositor, when viewed from below, is slightly retracted inward and two pairs of valvulae are distinguishable in it (Fig. 6C). The subgenital plate is shifted proximally, relative to the proctiger, and probably does not cover the ovipositor from below.

The female terminalia of *@@ala* are very different: the poorly distinguishable proctiger is very short, carries the poorly visible anus dorsally and does not cover the ovipositor from above, while the small triangular subgenital plate slightly covers the ovipositor from below (Fig. 5E–H). The ovipositor is short, pale, with two visible pairs of valvulae: slender inner and powerful outer (Fig. 6F). The outer valvulae bear basal cuticular folds (Fig. 6E, G, H), curved thick apical setae, lateral setae (Fig. 6G) and cover the inner valvulae laterally (Fig. 6H). This structure is more reminiscent of the female genitalia of modern whiteflies (Aleyrodidae; Weber, 1935).

The great difference in the ovipositor structure of *@@ala* and *Dingla* may indicate that they had different oviposition strategies.

### 4.7. Male terminalia

As in the case of females, the male terminalia of *@@ala* differ markedly from those of other genera of Miralidae. In *Dingla* and *Burmala*, the male terminalia consist of (1) a long, flattened, caudally directed proctiger, with a serrated ventral surface and long setae at the edges, (2) a short, flat, pentagonal subgenital plate, and (3) long, caudally directed parameres (Fig. 7C–E). The parameres are armed with claws directed towards the proctiger and slightly towards each other. The parameres are closely connected or fused and appear to move as a single structure (bent down together in *Dingla shagria*, PIN 5608/177a).

The male terminalia of *@@ala* include (1) a short, conical, caudally directed proctiger with a flattened dorsal side, several thick setae and an apical membranous structure near the anus, (2) long, massive, scoop-shaped, caudally directed subgenital plate, with several sclerotized grooves on the ventral side, and (3) short parameres, each bearing a sclerotized claw at the base and two thick apical setae (Fig. 7A, B).

The basic set of armature on miralid parameres includes a preapical sclerotized claw (or a row of thick setae) and an apical process bearing thick and slender setae.

The male terminalia of Miralidae are directed caudally, whereas in modern psyllids they are directed dorsally (Hodkinson and White, 1979), except for Togepsyllinae (Luo et al., 2017; Fig. 7F).

### 4.8. Compound wax pores

Many phloem-feeding hemipterans excrete honeydew and use waxy secretions to protect themselves from getting trapped in it (Ammar et al., 2013a, b). Females and nymphs of Psylloidea s.str. have a circumanal ring of wax pores to safely package honeydew droplets with a coat of wax. Psyllid females propel such wax-packed droplets away, and nymphs sometimes shed them together with exuviae. In contrast, the males, lacking a circumanal ring, secrete honeydew without wax and place it onto the plants (Douglas, 2006; Ammar et al., 2013a). Among extinct psyllomorphs, the circumanal ring has been found only in the female of *Amecephala pusilla* Drohojowska, 2020 (Liadopsyllidae; Drohojowska et al., 2020b).

Psyllids (mostly nymphs) may have wax pores on other parts of the body (Ammar et al., 2015), but such pores have not previously been observed in fossils. For the first time among Cretaceous psyllomorphs, we discovered wax pores on the ventral side of the pregenital abdomen in Miralidae: these are located on the ventral laterotergites, and each group of pores is surrounded by a ring-shaped sclerite bearing a circle of short thick setae (Fig. 8A–D). To refer to this set of structures we use the term “compound wax pore.” These compound wax pores are three-dimensional: the ring-shaped sclerite surrounds a prominent central process (Fig. 8A). The arrangement of the wax pores differs between sexes (recorded in both sexes of *Dingla* and female *@@ala* and probably applies to the entire family): three pairs of compound wax pores on abdominal segments 4–6 in females and on abdominal segments 3–5 in males. In modern psyllids, wax pores on the pregenital abdomen of adults have been reported only in *Togepsylla* (Psylloidea: Aphalaridae: Togepsyllinae), which have oval wax fields without ring sclerites on the sternites 4–6 in both sexes (Luo et al., 2017, Fig 8E). Thus, the structure of the wax pores of adult miralids is unique among both extant and extinct Psylloidea s.l.

Fields of wax pores (referred to as wax plates) on the abdominal sternites are characteristic of adult whiteflies, which apply waxy powder to the body and wings using legs (Weber, 1931, 1935; Navone, 1987; Byrne and Hadley, 1988). Sexual dimorphism in the position and number of these wax plates is common among Aleyrodidae (Shcherbakov, 2000). Their wax plates are generally subrectangular and not similar to the compound wax pores of Miralidae, although in the male giant whitefly *Udamoselis* Enderlein, 1909, with three pairs of wax plates on sternites 3–5, the anterior (largest) plates are oval and have a central patch (connected by a stripe of cuticle to the edge), reminiscent of the central process of compound wax pores in miralids (Martin, 2007, figs 1, 14).

The compound wax pores of Miralidae are even more similar to the compound wax pores of whitefly nymphs. These latter, located laterodorsally on the expanded abdominal tergites, are much larger than simple pores and have their own circular or semicircular sclerite and a central process (Wang et al., 2016). The structure of nymphal compound wax pores varies among whitefly taxa and is used as a diagnostic feature (Hodges and Evans, 2005). Among the known Miralidae, the compound wax pores are mostly uniform, only in the female of *@@ala* the pores of the sternite 6 are enlarged (Fig. 8C). The presence of ring-shaped sclerites in the compound wax pores of miralids indicates that their waxy secretions could have a specific shape, like that of whitefly nymphs.

Despite the similarity in the structure of the wax pores of whitefly nymphs and Miralidae, it is difficult to imagine what function such pores could have in flying and actively moving adult miralids. It seems unlikely that these pores, located far from the anus, were used to pack sticky excrement. Perhaps, Miralidae applied waxy secretions from their ventral abdominal pores to coat the host plant surface at the feeding site to prevent accumulation of sticky feces, and used their long hind legs equipped with rows of setae to transport and distribute the wax.

### 4.9. On the origin of whiteflies

Diverse primitive whiteflies are known from Cretaceous ambers (Drohojowska and Szwedo, 2011; Chen et al., 2022). The first two genera of this group were described from Lebanese amber (Schlee, 1970) and subsequently placed into the extinct subfamily Bernaeinae (Zherikhin, 1980; Shcherbakov, 2000; Shcherbakov et al., 2020). Two genera and three species of Bernaeinae were recorded as compression fossils from the Middle-Late Jurassic of Kazakhstan and China and the Early Cretaceous of Mongolia (Shcherbakov, 2000; Drohojowska et al., 2019). In these archaic Aleyrodidae, the forewing venation is more complete than in modern whiteflies, with R and M+CuA bifurcated in the basal wing third, RA oblique, and unforked M and CuA reaching the apical margin. This venation pattern is directly derivable from that of Liadopsyllidae s.str. by reduction of the M and CuA forks, but is completely dissimilar to that of Miralidae with their R and M+CuA forks shifted to the nodal level, and transverse RA and CuA2. Whiteflies are highly specialized, but in some respects more primitive than modern psyllids, and retain some features of their extinct psylloid (s.l.) ancestors.

## 5. Conclusions

Our findings show that in mid-Cretaceous Burmese amber, psyllomorph hemipterans were even more diverse than previously known. We describe a new genus and species of primitive jumping plant lice, *@@ala @@ioides* gen. et sp. nov., closely related to the genera *Mirala, Burmala*, and *Dingla*. These four genera form a peculiar group, which is easily recognized by the modified forewing venation, dorsoventrally flattened body and other features. Thus, the family-group taxa Miralinae and Dinglidae are synonymized as Miralidae stat. nov. Miralids are unique in that they have a very long, flexible, annulated 2nd labial segment and compound wax pores on the abdominal venter.

## CRediT authorship contribution statement

**Grigory Ivanov:** Conceptualization; Methodology; Investigation; Resources; Validation; Visualization; Writing – original draft; Writing – review & editing. **Dmitry Vorontsov:** Methodology; Investigation; Resources; Validation; Visualization. **Dmitry Shcherbakov:** Conceptualization; Investigation; Supervision; Validation; Writing – original draft; Writing – review & editing.

## Data availability

Data will be made available on request.

## Acknowledgements

We thank Roman Rakitov for extensive assistance in analyzing the unique morphology of miralids and obtaining SEM images, Alexey Bashkuev and Ilya Minaev for bringing the miralid fossils to our attention, Daniel Burckhardt and George Poinar for sending us photographs of *Mirala*, Vladimir Gnezdilov and Eugenia Labina for the loan of *Togepsylla* specimens, and Valery Mun for assistance with fluorescent microscopes. The fluorescence images were taken using the equipment of the Core Facility of the Institute of Developmental Biology, Russian Academy of Sciences. The work of D.D.V. was conducted under the IDB RAS Government basic research program in 2024 No. 0088-2024-0011.

